# Quantifying the Localization of Histological Staining Markers within the GI Epithelial Unit Axis: A Gastrointestinal Spatial Pathology Plugin for ImageJ

**DOI:** 10.64898/2026.05.28.728613

**Authors:** Arion Dey, Jared A. Weis, Victoria G. Weis

## Abstract

Histological analysis is crucial for understanding gastrointestinal (GI) tract homeostasis and disease pathophysiology. Various histological stains are commonly used in research settings for assessing development, disease pathogenesis, and therapeutic impacts. Specifically in the ordered architecture of the GI epithelium, current semi-quantitative analysis of histological staining relies heavily on manual scoring rubrics and often lacks robust spatial assessment. To address this gap, we developed an open-source ImageJ plugin, building on a closed-source predecessor, aimed at analyzing the spatial localization pattern of user-defined points-of-interest, such as positively stained cells, along the GI epithelial units. The plugin, developed using ImageJ 1.53.0 and Java programming language in Eclipse, interfaces with ImageJ and leverages Java libraries for data processing. The workflow involves uploading a microscopy image of the GI tissue of interest, determining base and top orientation landmarks of the GI epithelial units through manual identification, and annotating point-of-interest coordinates using ImageJ. The output includes centroid coordinates for each point-of-interest, the absolute distances of the points from the base and top landmarks, and the normalized distances of the points relative to the total height of the GI unit. The plugin generates histograms for displaying average point-of-interest distances along the GI unit axis. This information facilitates quantification of total points-of interest counts, analysis of height localization within the GI unit, and determination of average GI unit heights. The plugin serves as a crucial tool for robustly assessing various biological mechanisms within the GI tract, including EdU localization, migration distances, changes in cell type localization, and identification of new expression patterns along the GI unit axis. Overall, this open-source ImageJ plugin provides a semi-automated, user-friendly solution for leveraging important insights into spatial localization of tissue histology expression patterns within the GI structured architecture, with streamlined post-processing pipelines for robust large-scale analysis.

## Introduction

Histological staining is a mainstay of tissue assessment and is critical for understanding tissue homeostasis and disease. In the research setting, tissue histology is widely used to assess healthy development and homeostasis, disease initiation and progression, and impacts of experimental therapeutic approaches. This tissue analysis often includes histological stains and immunohistochemistry for the quantification of specific cell types or protein expression within the tissue architecture to identify potential changes in either the quantity or localization. Assessment of tissue staining is often qualitatively interpreted, with some semi-quantitative analysis typically performed using global total cell counts or coarse and time-consuming subjective manual scoring assessments of localization. Robust assessment of spatial dependence encoded within tissue staining patterns is therefore often lacking in these qualitative and semi-quantitative assessments. This shortcoming is especially relevant in the analysis of the gastrointestinal (GI) epithelium, which is typically arranged in a spatially ordered architecture with regular repeating epithelial units, such as in the crypt-villus axis or gastric gland in the small intestine and stomach, respectively. Therefore, there is a compelling need within the GI epithelial biology field for an effective tool to analyze how ‘stained’ markers of interest are expressed within the GI epithelial unit axis and that allows for large quantities of tissue to be quantified.

In recognition of the need for objective and quantitative spatially dependent assessment of GI tissue, several groups have developed analysis methods to capture and analyze staining patterns along the height of the GI epithelial unit axis. One analysis method trisected the epithelial unit and recorded the fractional representation of cells within each area to quantify cell movement, specifically by separating the unit by mesenchymal zones (Brown et al., 2007). Similarly, another group quantified Ki67 expression within the GI epithelial unit through assigning the unit a “s(surface), m(middle) and b(base))” (Lavery et al., 2014). There is also quantification done through the manual division of the unit into individual cells followed by sectioning it into different cell groups to generate a position-based histogram (Stange et al., 2013). Previously, we developed a closed-source MATLAB tool to analyze the location of staining markers of interest, with output histograms denoting more accurate staining marker-of-interest density along the spatial height of the GI epithelial unit (Caldwell et al., 2022; Meyer et al., 2020; Weis et al., 2021; Weis et al., 2017). We applied this MATLAB tool to robustly analyze staining patterns in stomach and intestinal tissue, providing important spatial localization context to cell types and protein expression patterns in GI tissue. However, despite the development of various spatial analysis techniques by individual research groups, there are no existing open-source tools to enable quantitative analysis of GI epithelial units, precluding a wider adoption of spatial quantitation of tissue staining within the GI epithelial biology research community.

In this work, we present a simple and user-friendly open-source plugin to enable spatial analysis of GI histological staining patterns to provide important context on the distribution of markers-of-interest within the ordered GI epithelial unit axis. We describe a streamlined workflow detailing the installation, operation, and output of the plugin, along with two separate case studies demonstrating the utility of spatial histological analysis in gastric and intestinal tissues. The first case study demonstrates the utility of Spatial Pathology Plugin for analyzing intestinal tissue in an experimental animal model of necrotizing enterocolitis (NEC), an intestinal disease that affects premature infants and poses a significant challenge in neonatal care. Recent research has shown the potential of a human placental-derived stem cell (hPSC) therapy to ameliorate intestinal damage associated with NEC (Weis et al., 2021). Findings highlight the potential of hPSC as a therapeutic strategy for mitigating NEC-induced intestinal damage and underline their utility as a research tool for better understanding neonatal repair mechanisms in the context of NEC (Weis et al., 2021)

In the second case study, we show the utility of Spatial Pathology Plugin for identifying localization of cell lineage tracing and metaplasia markers in the stomach mucosa. The development of metaplasia, a phenomenon where differentiated cell types transform in response to tissue injury, holds significant relevance as it a precursor to cancer. In the context of gastric metaplasia, the cell lineages present in the reparative mucosa after injury are diverse and include foveolar cells, proliferating cells, and spasmolytic polypeptide expressing metaplasia (SPEM) cells, which are pivotal in the metaplastic process. As detailed in recent work (Caldwell et al., 2022), two potential origins have been considered: zymogen-secreting chief cells in the stomach mucosa and isthmal progenitor cells. Thus, the localization of lineage-traced cells within the gastric gland yields significant insight into the cellular dynamics and potential roles of each of these cell types in metaplasia initiation and progression. The use of spatial mapping has shed light on the predominant origin of SPEM cells, emphasizing the role of mature chief cells in the metaplastic process after acute mucosal injury, which has implications for comprehending gastric metaplasia and its potential connection to cancer progression (Caldwell et al., 2022).

## Materials and Methods

An open-source ImageJ plugin to analyze spatial patterns in markers-of-interest identified in histological staining of GI tissues was developed based on a closed-source MATLAB tool previously developed by our group (Weis et al., 2021). To confirm workflow and accuracy of the new ImageJ plugin, re-analysis of histological tissue staining localization patterns within the GI epithelial unit axis was performed using the developed Spatial Pathology Plugin from two prior published works in intestinal (Weis et al., 2021) and gastric (Caldwell et al., 2022) tissues. The complete methods for tissue acquisition, staining, and data collection have been previously reported under these prior publications but is briefly explained here. For the intestinal tissue previous study, experimental NEC was induced in a rat pup model through formula feeding, hypoxia, and lipopolysaccharide (LPS) exposure in newborn Sprague-Dawley rat pups over the course of four days (Weis et al., 2021). Animals received intraperitoneal injections of either saline or hPSC to investigate the impact of cell-based treatment. The study focused on assessing macroscopic and histological intestinal damage, epithelial cell composition, and inflammatory marker expression in the ileum of these neonatal rat pups. For the gastric tissue previous study, gastric tissues were analyzed from mature mice with a novel chief cell-specific driver allele and treated with L635 to induce acute mucosal injury to assess the question of the role of mature chief cells as the predominant origin of SPEM cells during the metaplastic process after acute mucosal injury (Caldwell et al., 2022). The study focused on mouse models and histological staining techniques to analyze gastric intrinsic factor (GIF) expression, a stomach tissue-specific gene present in mature chief cells.

## Results

### Operation and Implementation

Spatial Pathology Plugin was developed using ImageJ version 1.72 and Java version 8 (Schneider et al., 2012). The code base uses both Fiji, a distribution of ImageJ, and legacy functions of ImageJ as well as other Java libraries. To use Spatial Pathology Plugin, the user must have Fiji installed to use Fiji’s native update site functionality to install the software. Spatial Pathology Plugin can be operated on any installation of Fiji. It can be installed by calling the update site from Fiji’s native updater client.

Spatial Pathology Plugin introduces several new forms and redefinitions of previous analysis techniques. A primary feature is the processing of data using user-defined spatial location-of-interest markers in relation to the GI epithelial unit axis coordinate reference frame. This allows for quantification of positively stained cells or immunomarkers of interest within the GI crypt/villus or gastric gland axis, for example. Spatial Pathology Plugin extends upon our previously developed custom-built closed-source MATLAB software that analyzed staining and quantified spatial localization from user annotated images in Photoshop, by allowing for a user’s workflow to be entirely contained within the open-source ImageJ environment.

The plugin is installed through Fiji’s update site and started within the ImageJ/Fiji environment plugins menu. To use the plugin, as shown in Figure 1, the user selects the desired microscopy image to analyze from available image formats supported by ImageJ/Fiji. Then, the user is prompted with their chosen “bin interval” to be used in a subsequent histogram analysis. The ‘bin interval’ denotes the increments of the GI epithelial unit axis in which normalized localization distances will be quantified for histogram representation. Next, the user is prompted to denote locations designating the base and the top of the GI epithelial unit axis. Manually designated region-of-interest (ROI) lines are drawn by the user using either the freehand or segmented line tool and saved to the image as an ROI selection. The length and position of each line are recorded for subsequent calculations and data output. The user is next prompted to designate ROI points that denote staining locations of interest (e.g. positively stained cells or marker staining locations). This multipoint selection is added to the image with point coordinates recorded as output data and used for subsequent calculations. User designation of the base and top lines is essential to define the local coordinate reference of the GI epithelial unit axis, with relative and absolute distances to the ROI marker points serving as the primary spatial analysis outcomes of this plugin. Following user-designation of all ROIs, the plugin performs background calculations for output metrics, including absolute distances from each ROI point to both the base and top of the GI epithelial unit, normalized distances of each ROI point designating relative height position within the GI epithelial unit axis, and length of the base and top lines denoting GI epithelial boundaries. Calculation of quantitative output metrics is further detailed below.

**Figure 1:**
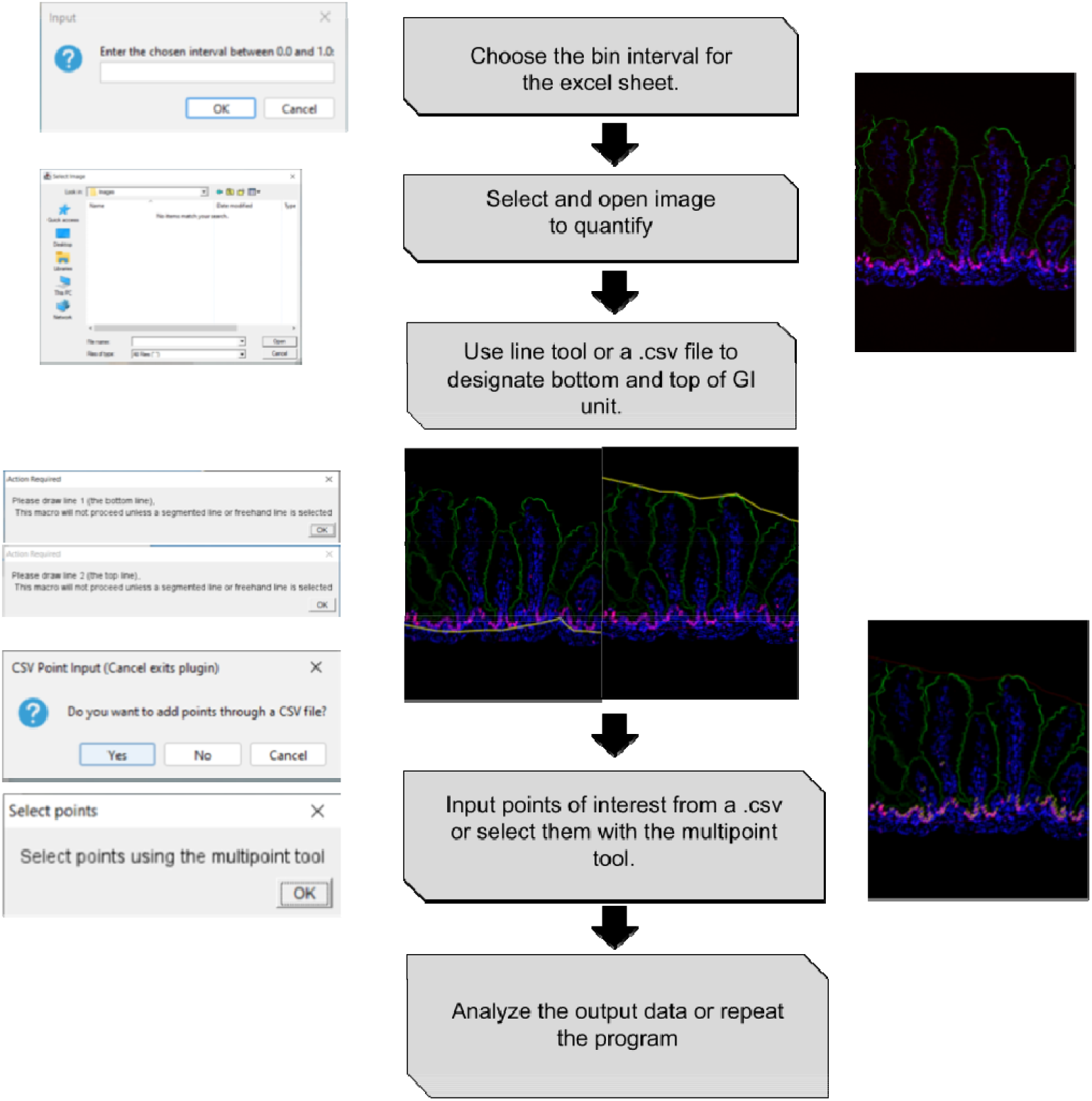
Schematic demonstrating the user workflow of Spatial Pathology Plugin with demonstrations and pictorial examples for the steps required for generating quantitative localization analysis. The steps required are: choose the bin interval for analysis, select and open the image to quantify, use the line tool or a .csv file to designate the bottom and top of the GI unit, input points of interest from a .csv file or select them with the multipoint tool, and analyze the output data or repeat the program.

### Output Metrics

The output includes several quantitative features that analyze the spatial relationship between the selected GI epithelial unit boundaries and staining marker points-of-interest. This comprehensive analysis is a fundamental aspect of understanding the spatial distribution of these points within the context of the GI epithelial unit. Measurements for each user-selected point from both the top and base of the GI epithelial unit are calculated for each point with respect to each line using an absolute distance formula. The Euclidean distance from point to line equation is as follows in Eq. 1:

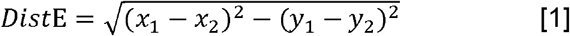

where *x*_1_ and *y*_1_ are coordinates on the top or base line and *x*_2_ and *y*_2_ are coordinates from a specific marker-of-interest point. Here, the locations of *x*_1_ and *y*_1_ are determined based on data extracted from ImageJ’s native ROI manager, ensuring accuracy in the spatial measurements. This calculation serves to quantify the exact spatial relationship between each point and the reference lines, providing precise measurements for further analysis. A normalized version of this distance metric is then calculated using Eq. 2:

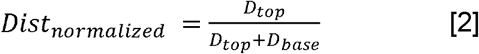

where *D*_*top*_ is the distance of the point from the top line and *D*_*base*_ is the distance of the point relative to the base. This offers a standardized representation of the spatial relationship for ease of comparison and analysis. The overall length of the base and top lines is also recorded, as calculated through a cumulative distance formula given by Eq. 3:

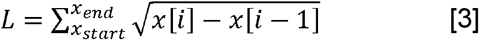

where *L* is the length of the line and *x* represents the x coordinate of points on the line and *i* represents a particular point on the line, *i*’s maximum is the full number of points on the line. This summation of distance based on each line’s point coordinates ensures a comprehensive measurement of each line’s total length, taking into account all relevant point coordinates. This detailed record is essential for a thorough spatial analysis and understanding of the relationship between the reference lines and the staining points-of-interest. This information is crucial in understanding the scale of the reference lines and their spatial significance in relation to the GI epithelial unit.

Lastly, the plugin also outputs a copy of the annotated image and a Microsoft Excel file containing the coordinates of the user-designated ROIs and the absolute and normalized distances of each point from each user-designated line. This detailed data export allows for further in-depth analysis and record-keeping, enhancing the utility of the plugin for research purposes. Following user annotation for all images within an analysis session, the plugin calculates and plots a histogram displaying the average point-of-interest spatial locations for the session. The histogram displays the spatial location distribution of points-of-interest within the GI epithelial unit coordinate system reference frame, with height locations on the y-axis denoting the average normalized distances from the base of the GI epithelial unit for each point. This spatial distribution data offers insights into the distribution of points of interest within the analyzed tissue, potentially revealing spatial patterns that might not be apparent through visual inspection alone. The normalization reflects ‘0’ as a point located at the base of the GI epithelial unit and ‘1’ as a point located at the top of the GI epithelial unit. This normalization simplifies the interpretation of the data, allowing for a consistent reference frame and making it easier to compare spatial data across multiple sessions or studies. The user selected bin increment denotes refinement in the relative location position in the x-axis of the histogram. This customization option allows researchers to adapt the analysis to their specific needs, enabling the precise exploration of spatial patterns within the GI epithelial unit and enhancing the accuracy of the findings.

### Case study: Localization of intestinal stem cell and proliferation/apoptosis markers in necrotizing enterocolitis in the neonatal intestine

As shown in Figure 2, Spatial Pathology Plugin can be used to analyze the spatial distribution of histological markers of interest in rat pup intestinal tissue in an experimental model of NEC disease. By visualizing the precise localization of critical cell populations within the crypt-villus axis of the neonatal intestine, including Paneth cells, SOX9+ cells, and LGR5+ stem cells, Spatial Pathology Plugin can offer valuable insights into the spatial dynamics of NEC disease pathogenesis and therapeutic regeneration response. This spatial analysis provides a more comprehensive understanding of cellular responses to disease progression and how therapy may promote epithelial healing and the repair of NEC-induced intestinal damage as compared to global cell counts. An example of this spatial analysis is shown in Figure 2B, demonstrating that cells marked with Ki67, a marker of proliferating cells, remain relatively close to the base within the intestinal crypt region. Similarly, OLFM4, a marker of LGR5 stem cells, shows a heavy concentration near the base of the crypt. In contrast, Ezrin, a marker of enterocytes, shows low density within the base, with somewhat uniform distribution within the rest of the villi. Combined, these spatial analysis tools offer additional contextual insight which is typically overlooked by raw global cell counts and can provide insights into cell- and molecular-level presentations of NEC.

**Figure 2:**
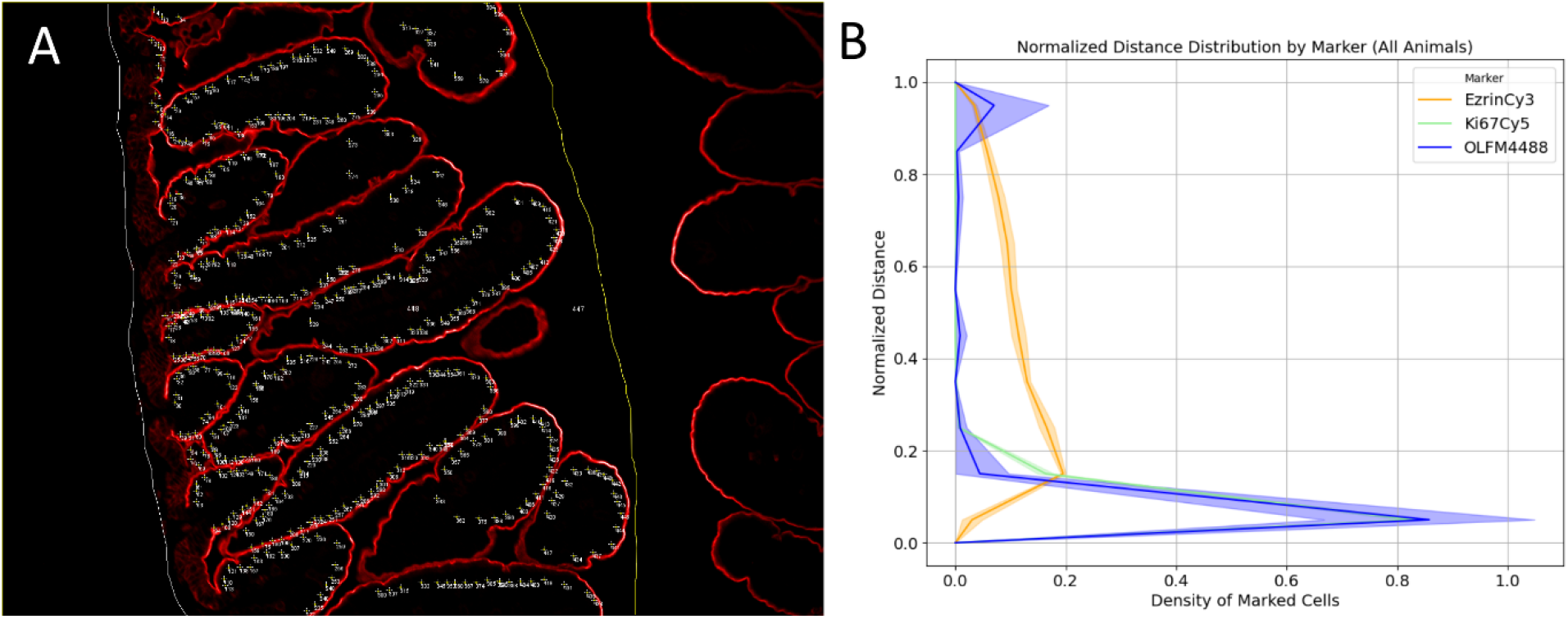
Spatial Pathology Plugin applied to analysis of intestinal data to quantify localization along the crypt-villus axis. (A) An example annotated image of a control (non-NEC) intestine marked with Ezrin. (B) Marker stratified distribution histogram using normalized distance data generated directly from Spatial Pathology Plugin.

### Case study: Localization of cell lineage tracing and metaplasia markers in the stomach mucosa

In order to further validate the utility of Spatial Pathology Plugin, we sought to replicate previous results using data from the stomach mucosa of GIF-GFP reporter mice who were administered Dox and treated with 1 dose of L635 by oral gavage for the purpose of triggering the metaplastic process (Caldwell et al., 2022). The distribution as indicated by utilizing Spatial Pathology Plugin, allows investigation into how different cell types are arranged along the gastric gland during both normal homeostasis and in response to acute mucosal injury, a pivotal trigger for metaplastic transformation. The plugin enables analysis of spatial niches within the GI epithelial unit where metaplasia is more likely to manifest, enabling in-depth study of precursor cell population localization. For comparison, the analysis was done across a population of metaplastic stomach mucosa and normal/nontreated stomach mucosa (Figure 3A). Similarly to previous findings, in normal gastric glands the GFP-labelled cells were shown to have a generalized regional restriction to the base of the gland, therefore, there were relatively few GFP-labelled cells that were both in the upper area of the gland and were copositive for GSII (Figure 3B). Whereas in the case of metaplastic glands, we find similarly to Caldwell et al. that comparatively there was a much higher rate of regional distribution to the top of glands. This confirmation of results further demonstrates the dynamic plasticity of SPEM cells under metaplastic conditions and underscores the utility of Spatial Pathology Plugin.

**Figure 3:**
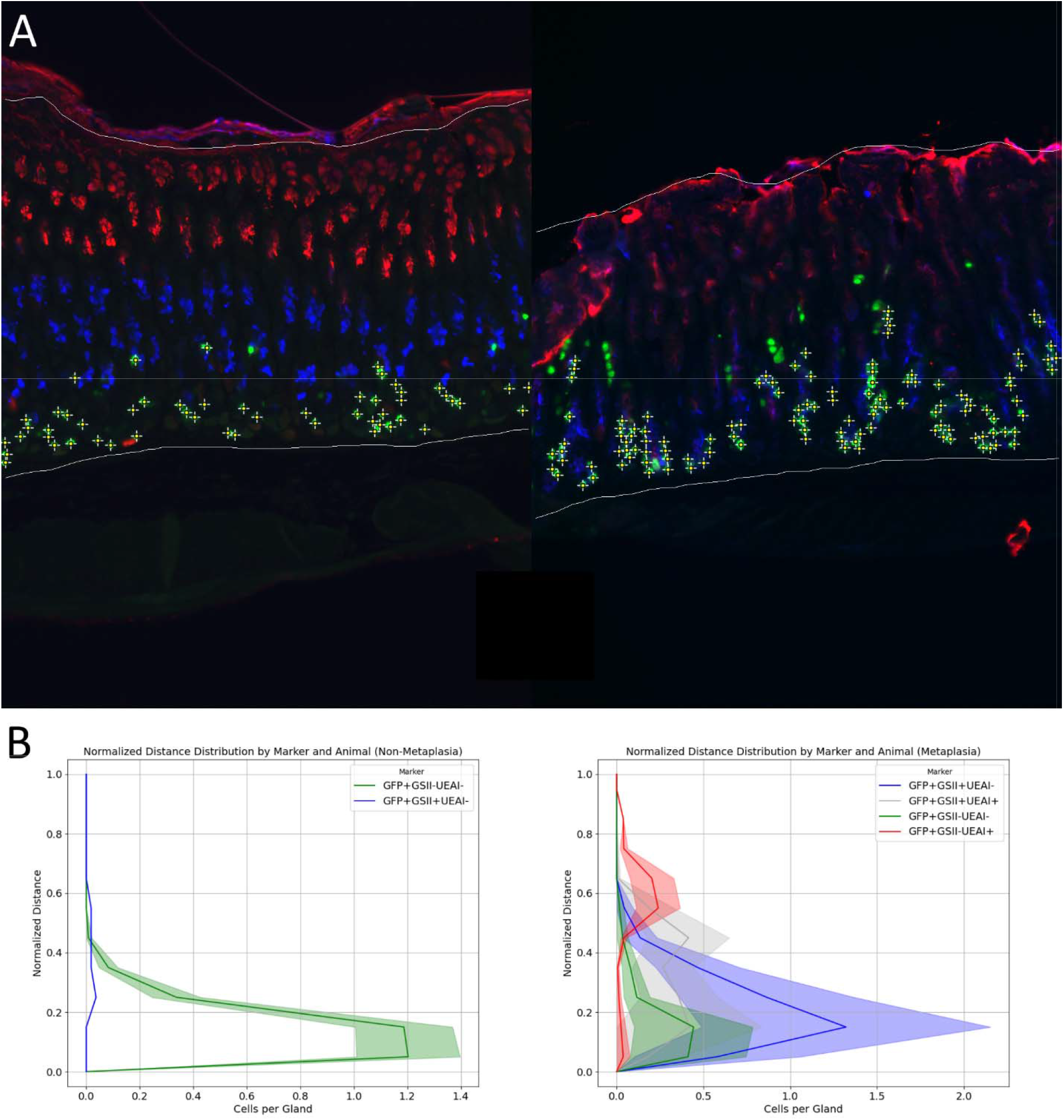
Spatial Pathology Plugin applied to analysis of metaplastic and normal stomach mucosa to quantify localization along the gastric gland. (A) Example images of normal stomach mucosa (left) and stomach mucosa that has undergone metaplastic process (right) with annotations performed using Spatial Pathology Plugin. (B) Marker stratified distribution histograms of a population of normal (left) and metaplastic (right) stomach mucosa using normalized distance data generated direclty from Spatial Pathology Plugin.

## Discussion

The open-source format and integration of Spatial Pathology Plugin within Fiji facilitates ease-of-use and accessibility within the research community. This plugin, hosted on Fiji’s update sites, is designed to be user-friendly and specialize in the analysis of histological staining of gastrointestinal (GI) tissues. The software’s capability to record spatial data is significant for various analytical applications, particularly in the context of research involving tissue morphology and cellular localization. Spatial Pathology Plugin is particularly valuable for studying the spatial localization of intestinal stem cells and proliferation/apoptosis markers in conditions such as necrotizing enterocolitis (NEC) in neonatal intestines. By providing detailed spatial localization data, the software aids in understanding the complex organization and interaction of cells within the small intestinal epithelium along the crypt-villus axis. This spatial analysis is crucial for exploring the developmental pathways and mechanisms of cell differentiation and proliferation, which are essential for unraveling complexities of tissue morphogenesis and advancing neonatal care. Moreover, Spatial Pathology Plugin enhances research capabilities by enabling in-depth spatial analysis of therapeutic responses, such as those observed in hPSC therapy for NEC. By visualizing cell distribution and interactions, the software provides insights into how hPSC therapy promotes epithelial healing and repairs NEC-induced intestinal damage. This advanced level of spatial analysis supports the identification of specific mechanisms through which therapeutic interventions exert their effects, contributing to the development of more effective treatments for NEC and other neonatal conditions. Due to the insights that spatial analysis provides to these cases, an analysis of the localization of markers used in the visualization within the case studies is used to demonstrate the overall utility of Spatial Pathology Plugin.

### Case study: Localization of intestinal stem cell and proliferation/apoptosis markers in necrotizing enterocolitis in the neonatal intestine

In the case of intestinal stem cell localization, the small intestinal epithelium is organized into villi and crypts with intestinal stem cells (ISCs). This arrangement guides progression of cells through cycles of proliferation and differentiation. The crypts contain two subtypes of ISCs: the highly proliferative crypt base columnar cells (CBCs) and the quiescent reserve ISCs located at +4 to +6 positions from the crypt base. The spatial interplay of these respective ISCs contributes to the generation of differentiated cells, reflected by the upward migration of differentiated cells along the crypt-villus axis, and with the downward movement of Paneth cells. While well studied in the adult intestine, the developmental pathways in the neonatal intestine are less understood. The foundational knowledge of these developmental pathways is essential for unraveling tissue morphogenesis complexities and exploring neonatal conditions such as necrotizing enterocolitis (Venkatraman et al., 2021). In this particular test case involving NEC and hPSC therapy, Spatial Pathology Plugin, specifically designed for spatial analysis within the GI epithelial unit axis, plays a crucial role in enhancing the depth of insights. Previous research focused on visualizing the precise localization of hPSC within the intestinal tissue and their impact on the restoration of critical cell populations such as Paneth cells, SOX9+ cells, and LGR5+ stem cells. Spatial Pathology Plugin enables researchers to delve deeper into the spatial dynamics of the therapeutic response. This in-depth spatial analysis offers a more comprehensive understanding of how hPSC therapy promotes epithelial healing and aids in the repair of NEC-induced intestinal damage. By providing a visual representation of cell distribution and interactions, Spatial Pathology Plugin enhances the ability to pinpoint mechanisms through which hPSC therapy exerts its beneficial effects, potentially paving the way for further advancements in the treatment of NEC and neonatal care.

### Case study: Localization of cell lineage tracing and metaplasia markers in the stomach mucosa

In this case involving the study of metaplasia in the context of gastric injury, Spatial Pathology Plugin, designed for spatial analysis within the GI epithelial unit axis, offers support to analyze the gastrointestinal gland. The process of metaplasia, where differentiated cell types transform in response to tissue injury, is a crucial area of study due to its relevance as a precursor to cancer. The cell lineages involved in reparative mucosa after injury, such as foveolar cells, proliferating cells, and spasmolytic polypeptide-expressing metaplasia (SPEM) cells, are integral to the metaplastic process. Prior investigations, using mouse models and specific staining techniques to examine gastric intrinsic factor (GIF) expression, have shed light on the primary origin of SPEM cells, due to the significance of mature chief cells in the metaplastic process following acute mucosal injury. This knowledge holds implications for understanding gastric metaplasia and its potential link to cancer progression (Caldwell et al., 2022). With Spatial Pathology Plugin, we can delve into how different cell types are spatially organized during both normal homeostasis and in response to acute mucosal injury, a pivotal trigger for metaplastic transformation. The plugin facilitates an analysis of spatial niches within the GI unit where metaplasia is more likely to manifest, enabling the study of precursor cell populations.

## Conclusions

The GI spatial pathology plugin is an analysis tool that enhances the assessment of various biological mechanisms within the GI tract. While our work highlights several specific use cases in stomach and intestinal tissue, it’s important to recognize that spatial histological analyses have much broader applications across many different GI epithelial contexts. The identification of quantitative distributions of staining points-of-interest relative to the GI epithelial unit axis reference frame offers essential context not only about the overall abundance of positively stained locations but also their precise height, both in absolute and relative terms, within the GI epithelial unit. This data further informs the understanding of the spatial characteristics of histological expression patterns. Moreover, the potential for further analysis by using the output data to generate histograms that display the average and standard deviation of point-of-interest counts at various locations along the epithelial unit axis, enable quantitative and statistical comparisons of spatial patterns in GI tissue staining. These histograms provide a means to statistically evaluate and comprehend the spatial distribution of staining points, yielding valuable insights into the biological processes under study. Spatial Pathology Plugin serves as a semi-automated, user-friendly tool that offers vital insights into the spatial localization of tissue histology expression patterns within the structured architecture of the GI tract. Additionally, it streamlines post-processing pipelines, making it particularly well-suited for large-scale, robust analysis of spatial histological data, thereby advancing the data analysis capabilities of researchers in the field.

## Acknowledgements

These studies were supported by NIH-NIDDK K01DK125633, R01DK135955, American Gastroenterological Association Research Scholar Award in Health Disparities, and Wake Forest University School of Medicine Faculty Start-Up Funds.

## Data and code availability

The ImageJ plugin is available at: https://github.com/ariondey/Spatial-Pathology-Plugin and further described at: https://imagej.net/plugins/spatial-pathology-plugin

## References

Belev, B. (2013). Role of Ki-67 as a prognostic factor in gastrointestinal stromal tumors. World Journal of Gastroenterology, 19(4), 523. 10.3748/wjg.v19.i4.523

Brown, S. L., Riehl, T. E., Walker, M. R., Geske, M. J., Doherty, J. M., Stenson, W. F., & Stappenbeck, T. S. (2007). Myd88-dependent positioning of Ptgs2-expressing stromal cells maintains colonic epithelial proliferation during injury. Journal of Clinical Investigation, 117(1), 258–269. 10.1172/JCI29159

Caldwell, B., Meyer, A. R., Weis, J. A., Engevik, A. C., & Choi, E. (2022). Chief cell plasticity is the origin of metaplasia following acute injury in the stomach mucosa. Gut, 71(6), 1068–1077. 10.1136/gutjnl-2021-325310

Lavery, D. L., Nicholson, A. M., Poulsom, R., Jeffery, R., Hussain, A., Gay, L. J., Jankowski, J. A., Zeki, S. S., Barr, H., Harrison, R., Going, J., Kadirkamanathan, S., Davis, P., Underwood, T., Novelli, M. R., Rodriguez-Justo, M., Shepherd, N., Jansen, M., Wright, N. A., & McDonald, S. A. C. (2014). The stem cell organisation, and the proliferative and gene expression profile of Barrett’s epithelium, replicates pyloric-type gastric glands. Gut, 63(12), 1854–1863. 10.1136/gutjnl-2013-306508

Schneider, C. A., Rasband, W. S., & Eliceiri, K. W. (2012). NIH Image to ImageJ: 25 years of image analysis. Nature Methods, 9(7), 671–675. 10.1038/nmeth.2089

Stange, D. E., Koo, B.-K., Huch, M., Sibbel, G., Basak, O., Lyubimova, A., Kujala, P., Bartfeld, S., Koster, J., Geahlen, J. H., Peters, P. J., van Es, J. H., van de Wetering, M., Mills, J. C., & Clevers, H. (2013). Differentiated Troy+ Chief Cells Act as Reserve Stem Cells to Generate All Lineages of the Stomach Epithelium. Cell, 155(2), 357–368. 10.1016/j.cell.2013.09.008

Venkatraman, A., Yu, W., Nitkin, C., & Sampath, V. (2021). Intestinal Stem Cell Development in the Neonatal Gut: Pathways Regulating Development and Relevance to Necrotizing Enterocolitis. Cells, 10(2), 312. 10.3390/cells10020312

Weis, V. G., Deal, A. C., Mekkey, G., Clouse, C., Gaffley, M., Whitaker, E., Peeler, C. B., Weis, J. A., Schwartz, M. Z., & Atala, A. (2021). Human placental-derived stem cell therapy ameliorates experimental necrotizing enterocolitis. American Journal of Physiology-Gastrointestinal and Liver Physiology, 320(4), G658–G674. 10.1152/ajpgi.00369.2020

